# No evidence of evolution of male fitness components in response to yeast deprivation: Putative female specific nutrient dependent reproductive adaptation

**DOI:** 10.64898/2026.11.23.768529

**Authors:** Smita Adhikary, Rabi Sankar Pal, Subhasish Haldar, Bodhisatta Nandy

**Affiliations:** 1Department of Biological Sciences, Indian Institute of Science Education and Research Berhampur, At/Po: -Laudigam, Dist.- Ganjam, Brahmapur, Odisha 760003, India; School of Arts and Sciences, Division of Biological and Life Sciences, Ahmedabad University, Ahmedabad 380009, Gujarat, India; Mallick Para, Krishnagar, Nadia, West Bengal 741101, India

**Author notes:** Corresponding author, contact number.

**Keywords:** Intralocus sexual conflict, reproductive adaptation, protein deprivation, sperm defense, sperm offence, sex-specific adaptation

## Abstract

Intersexual genetic correlation (IGC) is pervasive in sexually reproducing organisms and can constrain sex-specific adaptive evolution. Nevertheless, sexes frequently undergo adaptive divergence in response to novel ecological challenges. Here, we investigated whether adaptation to nutritional stress is sex-limited or constrained by shared genetic architecture. We utilized four laboratory-adapted populations of *Drosophila melanogaster* subjected to experimental evolution under adult-stage live-yeast deprivation for over 140 generations—a regime that previously drove adaptive female reproductive evolution, particularly enhancing early-life fecundity and ovarian function. Because female *D. melanogaster* rely heavily on dietary yeast for protein-dependent egg production, this evolutionary response may be female-limited. Alternatively, male fitness traits could respond either through direct selection or via correlated response driven by IGC. To distinguish between these hypotheses, we evaluated a comprehensive suite of pre- and post-copulatory male competitive fitness components. We found that all measured male fitness traits remained unchanged following experimental evolution, demonstrating that nutrient-dependent adaptation in these populations is strictly female-specific, with no evidence of correlated responses or sexually antagonistic trade-offs. Our results indicate that the genetic architecture underlying nutrient-dependent reproductive adaptation is largely decoupled between the sexes, allowing each sex to independently approach its sex-specific fitness optimum.

## Introduction

Sexes often differ in strategies to maximize Darwinian fitness due to inherent differences in reproductive physiology, breeding ecology, and parental investment, leading to sexual conflict (Bateman, 1948; Chapman et al., 2003). However, because the same genome is expressed in both sexes, the evolution of life history, behavioural, and physiological traits in one sex can depend on their expression in the other—a phenomenon driven by intersexual genetic correlation (IGC; Lande, 1980; Bonduriansky & Chenoweth, 2009; Pennell & Morrow, 2013). Consequently, selection acting on one sex can induce a correlated response in the trait values of the other sex. The magnitude of IGC is typically quantified as the correlation between the additive genetic effects of both sexes for the expression of a given trait (Lande, 1980; Bonduriansky & Chenoweth, 2009; Poissant et al., 2010; Collet et al., 2016).

The role of IGC in constraining adaptive evolution has been demonstrated in numerous theoretical (Lande, 1980; Connallon & Hall, 2016) and empirical studies (Chippindale et al., 2001; Rice et al., 2006; Brommer et al., 2007). In a meta-analysis of 114 studies across diverse taxa, Poissant et al. (2010) evaluated IGC as a potential constraint on the emergence of sexual dimorphism. They found that positive IGCs predominate for morphological, behavioural, and developmental traits, suggesting that a shared genetic architecture can generate an evolutionary tug-of-war that limits the divergence of sexual dimorphism. In contrast, fitness components often exhibit low or negative IGCs, suggesting that the genetic architecture underlying fitness-related traits between the sexes may be weakly correlated, thereby offering greater scope for sex-specific evolutionary responses (Brommer et al., 2007). Nevertheless, sexually antagonistic selection—where an allele beneficial to one sex is detrimental to the other—can constrain sex-specific adaptation (Chippindale et al., 2001; Brommer et al., 2007). The impact of such constraints becomes particularly pronounced under stressful environmental conditions, where selection pressures differ substantially between the sexes (Arnqvist & Rowe, 2005; Perry & Rowe, 2018).

Nutrient-dependent evolution presents a compelling context in which the role of IGC remains understudied, despite clear scope for both sex-specific selection and shared genetics. Because reproductive investments differ between sexes, response to nutritional challenges (in terms of either quality or quantity) is also sex-specific (Ng et al., 2019; Brüggemann et al., 2025).

Certain forms of dietary restriction result in lifespan extension, likely due to a re-allocation of resources away from reproduction and toward somatic maintenance (Piper et al., 2011; Simpson et al., 2017). Empirical evidence indicates that this lifespan-extending effect of nutritional deprivation is sex-biased, often being more pronounced in females than in males (Magwere et al., 2004; Nakagawa et al., 2012). Furthermore, studies employing the geometric framework for nutrition have revealed that macronutrient balances differentially affect survival and reproductive traits in the two sexes (Maklakov et al., 2008; Rapkin et al., 2017; Hawkes et al., 2021). For instance, *Drosophila melanogaster* females are generally more sensitive to alterations in dietary protein content, whereas males are more responsive to carbohydrate variations (Lee et al., 2008; Jensen et al., 2015). Similarly, Han and Dingemanse (2017) demonstrated sex-specific consequences of protein deprivation in the southern field cricket (*Gryllus bimaculatus*), where protein restriction significantly reduced male survival while female survival remained unaffected.

Despite these sex differences in nutrient dependence, nutrient signalling, processing, and metabolism share substantial physiological commonality between the sexes. The underlying genes involved in nutrient acquisition, signalling, and metabolism are largely shared. Highly conserved nutrient-sensing pathways, such as the insulin/insulin-like growth factor signalling (IIS) and target of rapamycin (TOR) pathways, regulate growth, metabolism, and reproduction in both sexes (Katewa & Kapahi, 2011). However, female fecundity is more strongly coupled to dietary protein availability via IIS/TOR signalling than male reproductive output, as IIS/TOR directly regulates oogenesis and vitellogenesis in response to amino acid availability (Drummond-Barbosa & Spradling, 2001; Nässel & Vanden Broeck, 2016).

Similarly, genes encoding nutrient transporters and metabolic enzymes are expressed in both sexes despite having sex-specific fitness impacts (Innocenti & Morrow, 2010; Mank, 2017; Camus et al., 2019). Consequently, strong nutrient-dependent selection acting on one sex could drive a correlated evolutionary response in the other, which may be adaptive, detrimental, or neutral depending on the underlying IGC.

In this study, we utilized a set of laboratory-adapted *Drosophila melanogaster* populations subjected to experimental evolution for over 140 generations under adult-stage (live) yeast deprivation. Live yeast serves as a crucial protein source for females, and its absence severely restricts female fecundity (Becher et al., 2012; Mirth et al., 2019). Thus, this regime imposed strong selective pressure on females while having minimal direct impact on males (Dasgupta et al., 2022). Dasgupta et al. (2022) demonstrated adaptive female reproductive evolution in these populations, characterized by higher early-life fecundity and steeper reproductive senescence compared to ancestral controls, driven by an elevated rate of oocyte production.

Here, we evaluated whether these populations exhibited a direct or correlated evolutionary response in male fitness-related traits. We tested three competing hypotheses: (a) Direct selection / Correlated adaptation: If males also experienced selection due to yeast deprivation (or if selection on females drove a concordant correlated response), evolved males will show adaptive improvements in one or more fitness components relative to ancestral controls. (b)

Intralocus sexual conflict: If nutrient-dependent female adaptation involved loci under intralocus sexual conflict, evolved males will show a decrement in fitness components (a trade-off). (c) Sex-limited evolution / Decoupling: If female adaptation was strictly sex-limited due to low or absent intersexual genetic correlation, male fitness components will show no response.

To test these hypotheses, we comprehensively evaluated pre- and post-copulatory male competitive fitness components in the evolved populations (YLBs) relative to their ancestral baseline populations (BLs; see Figure S1 for a complete schematic of evaluated traits).

## Methods

We have used four replicate populations of Yeast-less BLs - YLB_(1-4)_ that are selected for adult-stage yeast deprivation over more than 140 generations along with their respective four ancestral populations - BLs_(1-4)_ (Dasgupta et al., 2022). BLs are the laboratory-adapted outbred populations of *Drosophila melanogaster* maintained at a temperature of 25°C (±1°C) and 24-hour light condition on a 14-day discrete generation cycle with a population size of approximately 2800 individuals on standard banana-barley-jaggery-yeast food. The experimental YLB populations were derived from the control populations, and evolved under identical handling, except for the step wherein the BL populations are given *ad-lib* live-yeast supplement on day 12 of the generation cycle. Except for this, all other aspects of laboratory ecology and handling were kept identical between BL and YLB. We used another outbred population BL_st._ with a similar genetic background to BLs and having an autosomal recessive scarlet eye-colour (Nandy et al., 2016). Flies from this population were used in assays that required paternity measurement. All these experiments were performed in four statistical blocks, with each block corresponding to each replicate BL-YLB population set.

The experimental flies were generated following one generation of common garden rearing for both population sets, during which both control and experimental populations were grown under a condition identical to that followed for the control populations. All the experimental flies were generated under standard conditions (25°C, 60–70% RH, 24-hour light condition, and larval density ∼70 per 8ml of standard banana-barley-jaggery food per vial). All the flies used across the experiments were virgin (unmated) flies. Virgin flies were generated by collecting young flies within 6 hours of eclosion on light carbon dioxide anaesthesia, and were held until the assay in sex-segregated groups of 20 flies per vial with ample food. Three assays were performed to quantify various components of pre- and post-copulatory fitness traits in males. All the assays were conducted in glass vials (25 mm diameter × 90 mm height) with 3-4 mL of food.

### Experiment 1: Mating behaviour in presence of a competitor

Since a key component of male fitness is mate acquisition ability, a mating assay was set up to investigate three components of mating behaviour – success in securing a mate under competitive condition (mating success), time to initiate mating with a virgin female (mating latency), and duration of copulation as a measure of male investment per mating (copulation duration). To this effect, mating observation vials were set up with a target male (either BL or YLB), a competitor male and a virgin female. The assay female and the competitor male in such a setup were from the BL_st_ population. All flies in a mating vial were about 2days old virgin. Mating vials were standard culture vials with ample food seeded with a limiting quantity of yeast suspension (concentration = 11.25mg per mL per fly) to stimulate mating and mimic the conditions of the control population, where typically flies of this age are transferred to a population cage with *ad-lib* yeast supplement. 70 mating observation vials were set up per population. To set up a mating trial, adults were introduced in quick succession, and immediately transferred to an observer and left undisturbed. Every vial was observed until the completion of mating. Identity of the successful male, time of onset of mating, and time of end of mating were recorded.

### Experiment 2: Sperm defense assay

Another key component of male fitness is the ability to secure paternity after the female remates – i.e., under sperm competition. A double-mating assay was therefore set up to measure the sperm defense ability. It is often measured as the proportion of progeny sired by the first male (referred to as P1) in a double mating experimental setup. Along with P1, four other components of male fitness are usually considered important. (1) Time taken to initiate mating with an unmated female – mating latency, which is often considered a measure of mating ability or attractiveness of a male. (2) Duration for which a male copulates with a female – copulation duration. It may reflect a male’s acceptance by a female and is known to be a proxy for the amount of sperm and seminal fluid transfer, directly impacting fertilization success and paternity. (3) Mating fidelity is measured as remating delay of a male’s mate (remating latency) and the duration of remating (remating duration) when it actually happens. Since mating rate in *D. melanogaster* females is fairly high and egg production/oviposition is continuous, once a male has secured a mating, delayed remating by a female significantly improves the male’s paternity share. When the female actually remates, the longer the duration, the better would be the paternity share of the new mate. Hence, both aspects of remating behaviour are expected to be key determinants of male fitness.

We measured mating latency, copulation duration, remating latency, and remating duration using a sequential double mating design. Furthermore, the progeny produced by the female were scored for paternity share to measure P1. We used an autosomal recessive marker to do the latter.

In this assay, a mating observation vial was set up with a virgin female and a target male from the experimental populations, followed by remating with a second male from a standard stock. The assay female and the second male were from the BL_st_ population (see above).

Similar to the previous assay, all flies used were about 2-day old virgins. Mating observation vials were standard culture vials with ample food seeded with a limiting quantity of yeast suspension. 100 such mating observation vials were set up per population. After the completion of the first mating, males were discarded, and females were retained in the same vial. After 24 hours of the first mating, females were given a second mating opportunity (i.e., remating opportunity) by the introduction of two BL_st_ males. Upon introduction of these males, vials were left undisturbed and observed continuously for 6-hour. Within this 6-hour window, if a female was found to have initiated mating, the same was allowed to complete. Males were discarded immediately after the completion of mating. Females were retained in the same vials until oviposition (see later). Females that did not remate within this time window were given a second round of a similar 6-hour remating opportunity window on the following day, with continuous observation similar to that done on the first round.

After the second remating opportunity, all remated females were transferred individually to oviposition tubes (16mm diameter x 84 mm height with ample quantity of standard food) and were allowed to lay eggs for 18 hours. Females were then discarded, and the tubes were left undisturbed for the progeny to grow. On the 12^th^ day post oviposition, the tubes were frozen at -20 °C, and the progeny were scored for eye colour. Note that since the females and the second male were both from BL_st_, progeny sired by the second male were expected to be scarlet-eyed, whereas those sired by the first male were expected to be red-eyed. P1 was calculated for each remated female as the proportion of red-eyed progeny.

### Experiment 3: Sperm offence assay

In addition to the defense component of sperm competitive ability, the other key component of male fitness is sperm offense ability – viz., paternity success from mating with already mated females. Similar to P1, sperm offense ability is measured as P2 - proportion of offspring sired by a male that was the second mate of a female subjected to double-mating. In addition, time taken to initiate mating with an already mated female (remating latency) and duration of such matings (remating duration) may also be considered key male fitness components.

We conducted a sperm offense assay using BL_st_ female, BL_st_ male (first mate), and the target males (second mate) from the experimental populations. The assay followed the a similar general procedure, and sample sizes were identical to that of the sperm defense assay described earlier. Virgin BL_st_ females were first mated with the BL_st_ males, followed by remating with the target males after two days. Vials were manually observed to ensure that all females were mated. Once mating was completed, males were discarded and the females were retained in the same vial. 48 hours following the first mating, females were transferred to fresh vials, and target males were introduced. These vials were continuously observed for 8 hours by human observers for mating, and the time of onset and end of all matings were recorded. Time to remating (remating latency), and duration of for which each remating lasted (remating duration) were computed from these observations. Females that remated were transferred to oviposition tubes and allowed to lay eggs for 18 hours. 12 days post- oviposition, progenies were frozen at -20 °C, and the progenies were scored for eye colour. P2 was calculated for each remated female as the proportion of progeny that were wildtype red-eyed – i.e., sired by the second male.

### Data analysis

All the data were analysed using R software (version 4.2.2). In all models, male selection regime (levels: BL and YLB) was treated as a fixed factor, whereas block and all interactions that included block were treated as random factors. The interquartile range method was used to identify and remove the outliers (except for remating latency in Sperm defense assay). The Shapiro Wilk test was performed to check the normality of model residuals. LMM (linear-mixed model) was used when residuals were normally distributed. In cases where the normality test showed significant non-normality, we used GLMM (generalised linear mixed model). All tests were done using `lme4` package (Bates et al., 2015) and `glmmTMB` package (Brooks et. al., 2017) . The models for all the measured components of male fitness were listed in Table S3. Model fit was assessed using `DHARMa` package (Hartig, 2022). As the P1 and P2 data were found to be overdispersed, they were analysed using GLMM with the beta-binomial error distribution in the `glmmTMB` package (Brooks et. al., 2017). The `anova` function from the `lmerTest` package (Kuznetsova, 2017) was used to assess the effect of random factors through sequential model reduction. The `Anova` function from the car package (Fox & Weisberg, 2019) was used to obtain the p-values for the fixed effect predictor variables.

## Results

### Experiment 1

The effect of selection regime was found to be non-significant on mating latency, copulation duration, and mating success (Table1, Figure 1). Though there was a significant random effect of Block on mating latency and copulation duration, the effect of block×male selection regime interaction was found to be non-significant in all these analyses (Table S4).

**Figure 1:**
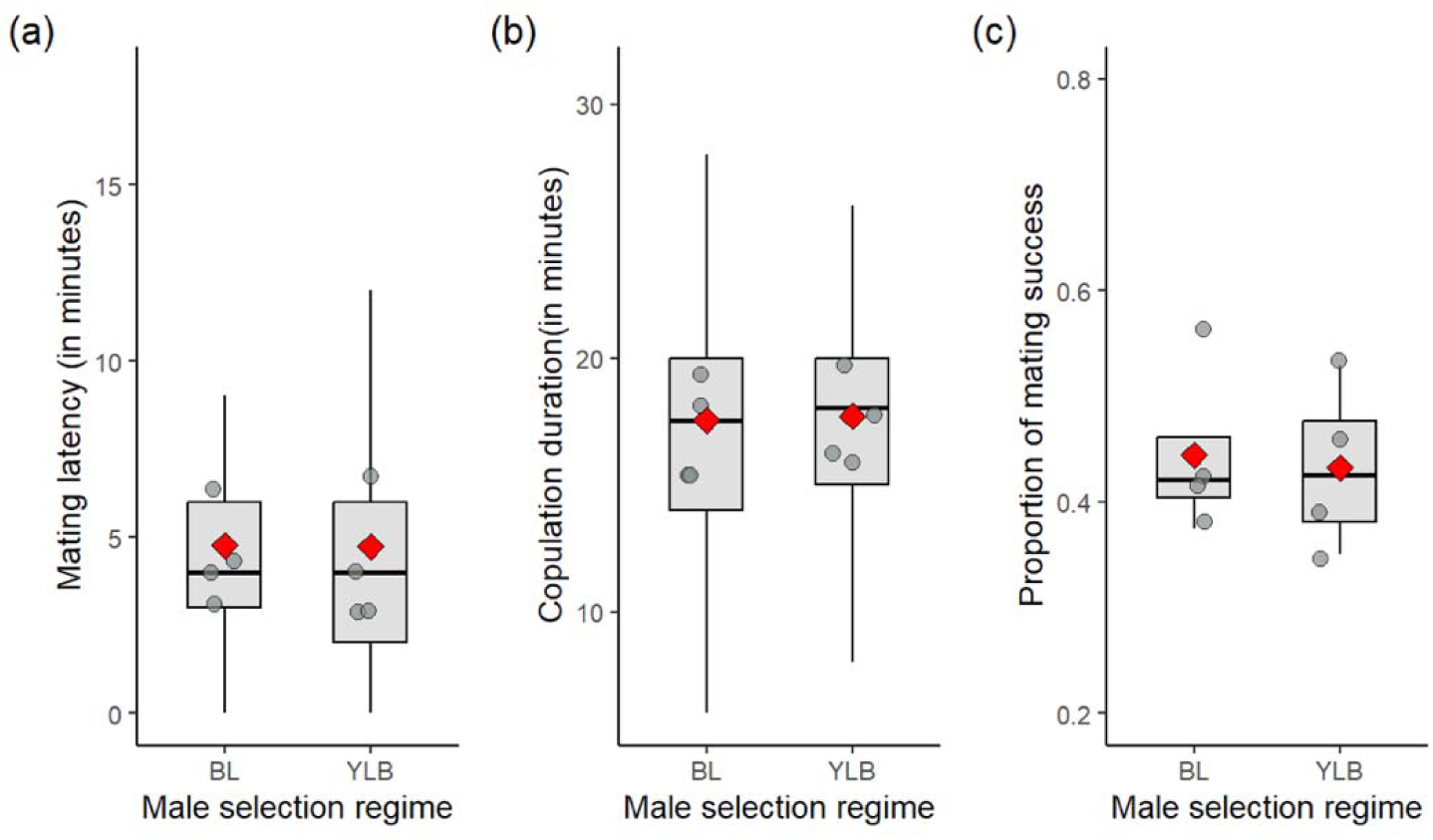
Boxplots showing (a) mating latency, (b) copulation duration and (c) mating success in experiment 1 (mating behaviour). The grey dots represent the mean values of the replicate blocks and red dot represents to the overall mean across replicate blocks.

### Experiment 2

The effect of selection regime was found to be non-significant on remating latency, remating duration, and P1 (Table 1, Figure 2). The effect of block×male selection regime interaction was not significant in any of these parameters (Table S5).

**Figure 2:**
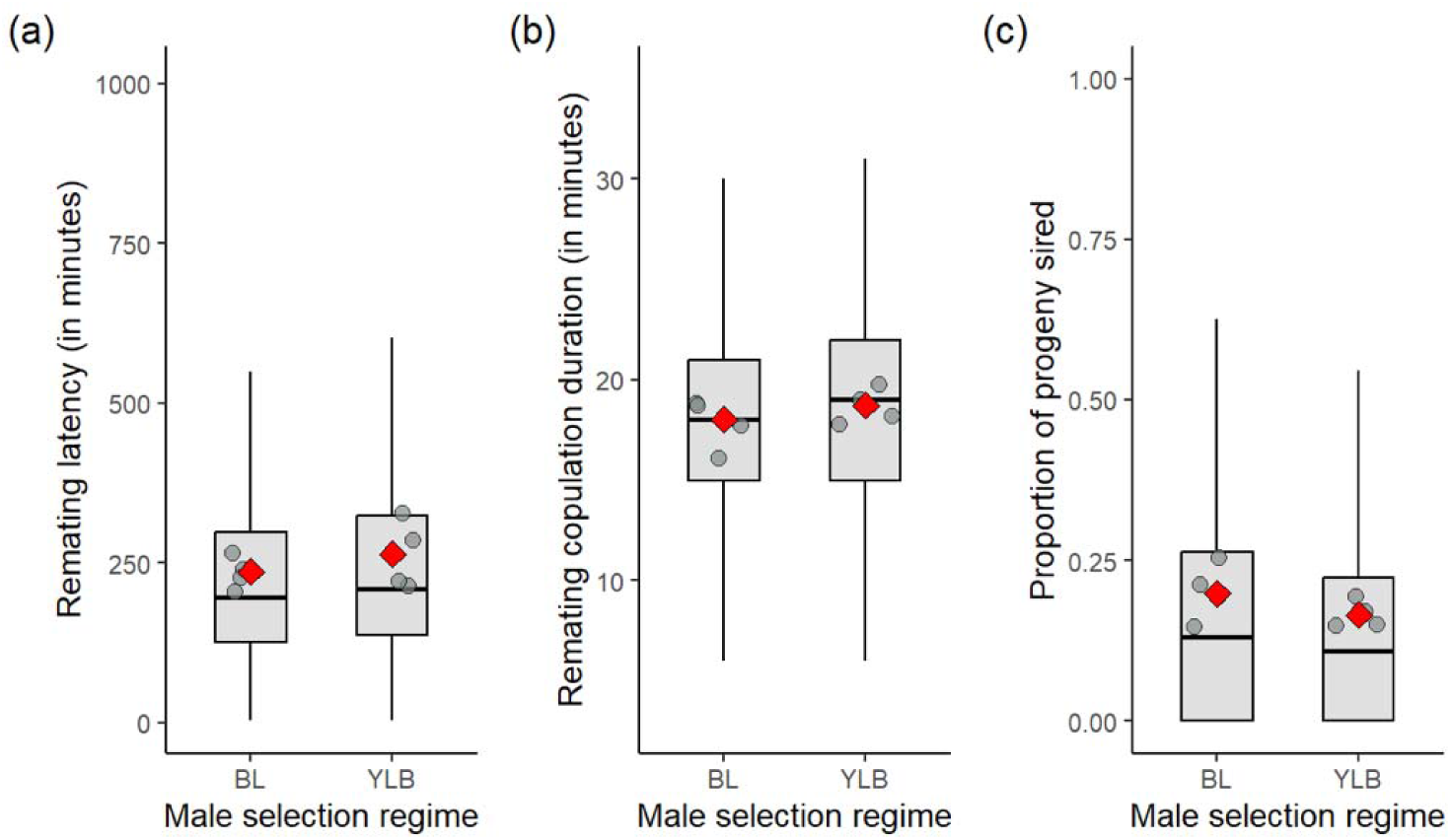
Boxplots showing (a) Remating latency, (b) Remating copulation duration and (c) P1 (proportion of progeny sired by target males) in experiment 2 (sperm defense assay). The grey dots represent the mean values of the replicate blocks and red dot represents to the overall mean across replicate blocks.

**Table 1:**
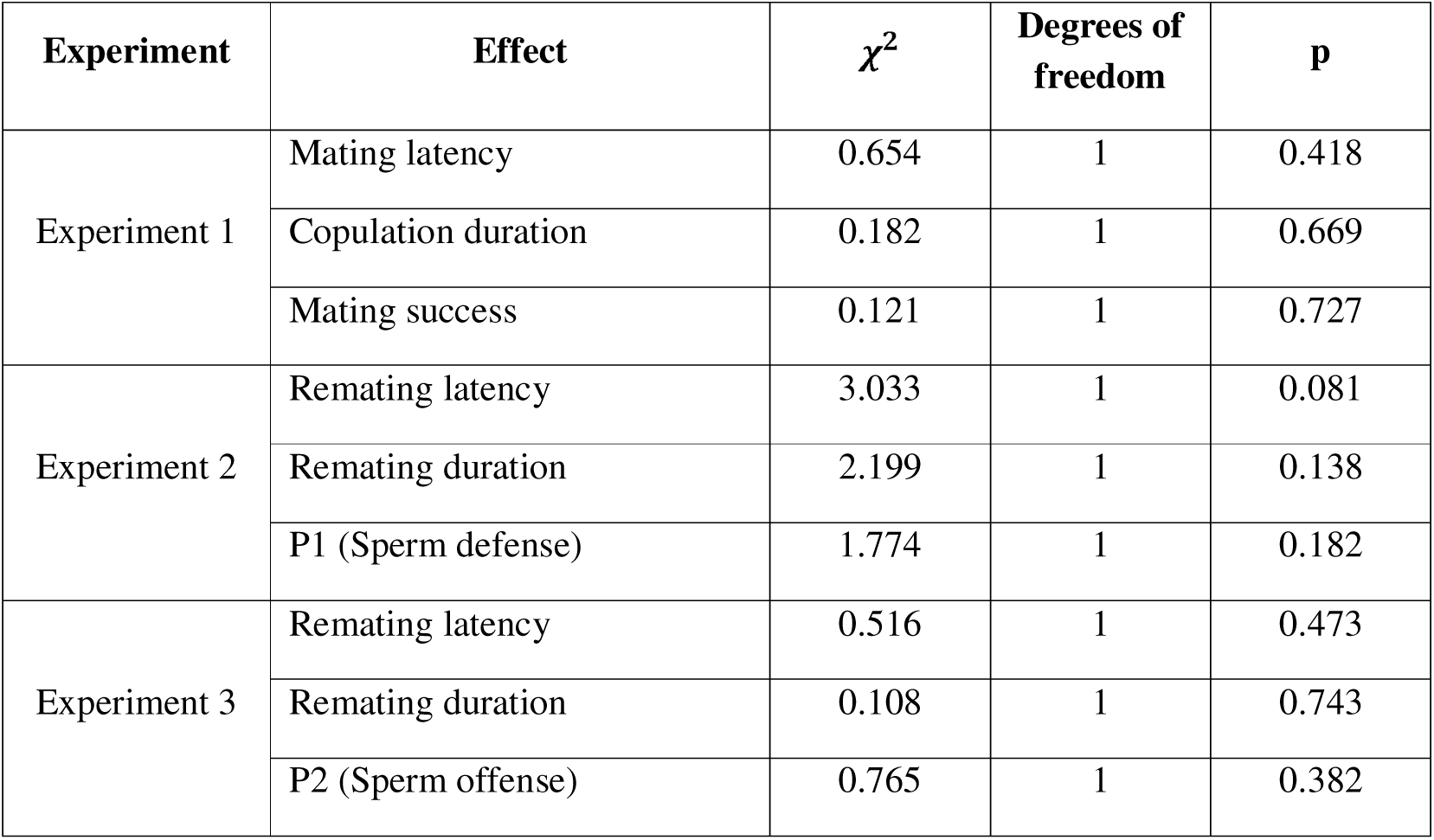
Results of the analyses of data from the three experiments using GLMM. The main effects of the fixed factors are shown. The exact models and the output of the analyses on the random factors can be found in the supplementary information file.

| **Experiment** | **Effect** | *χ*<sup>2</sup> | **Degrees of freedom** | **p** |
|---|---|---|---|---|
| Experiment 1 | Mating latency | 0.654 | 1 | 0.418 |
|  | Copulation duration | 0.182 | 1 | 0.669 |
|  | Mating success | 0.121 | 1 | 0.727 |
| Experiment 2 | Remating latency | 3.033 | 1 | 0.081 |
|  | Remating duration | 2.199 | 1 | 0.138 |
|  | P1 (Sperm defense) | 1.774 | 1 | 0.182 |
| Experiment 3 | Remating latency | 0.516 | 1 | 0.473 |
|  | Remating duration | 0.108 | 1 | 0.743 |
|  | P2 (Sperm offense) | 0.765 | 1 | 0.382 |

The effect of selection regime×block interaction was found to be significant on mating latency and copulation duration (Table S5), prompting us to carry out block-wise analyses of these datasets. Block-wise analysis of mating latency (Table S2, Figure S2) showed a significant effect of selection regime in the three of the blocks, while block 3 showed non-significant effect of selection regime. We therefore re-analyzed the mating latency results excluding this block from the analysis, and found that selection regime was now a highly significant predictor (Figure S3). YLB males were found to have ∼18% shorter mating latency. In case of copulation duration, we found significant effect of selection regime in two blocks – viz., block 2 and block 4 (Table S3, Figure S4), albeit in opposite directions. Hence, these significant effects of fixed factor can only be considered spurious.

### Experiment 3

The effect of selection regime was found to be non-significant on P2, remating latency, and remating copulation duration (Table 1, Figure 3). Though there was a significant effect of Block on remating copulation duration, none of these analyses indicated a significant effect of selection regime×block interaction (Table S6).

**Figure 3:**
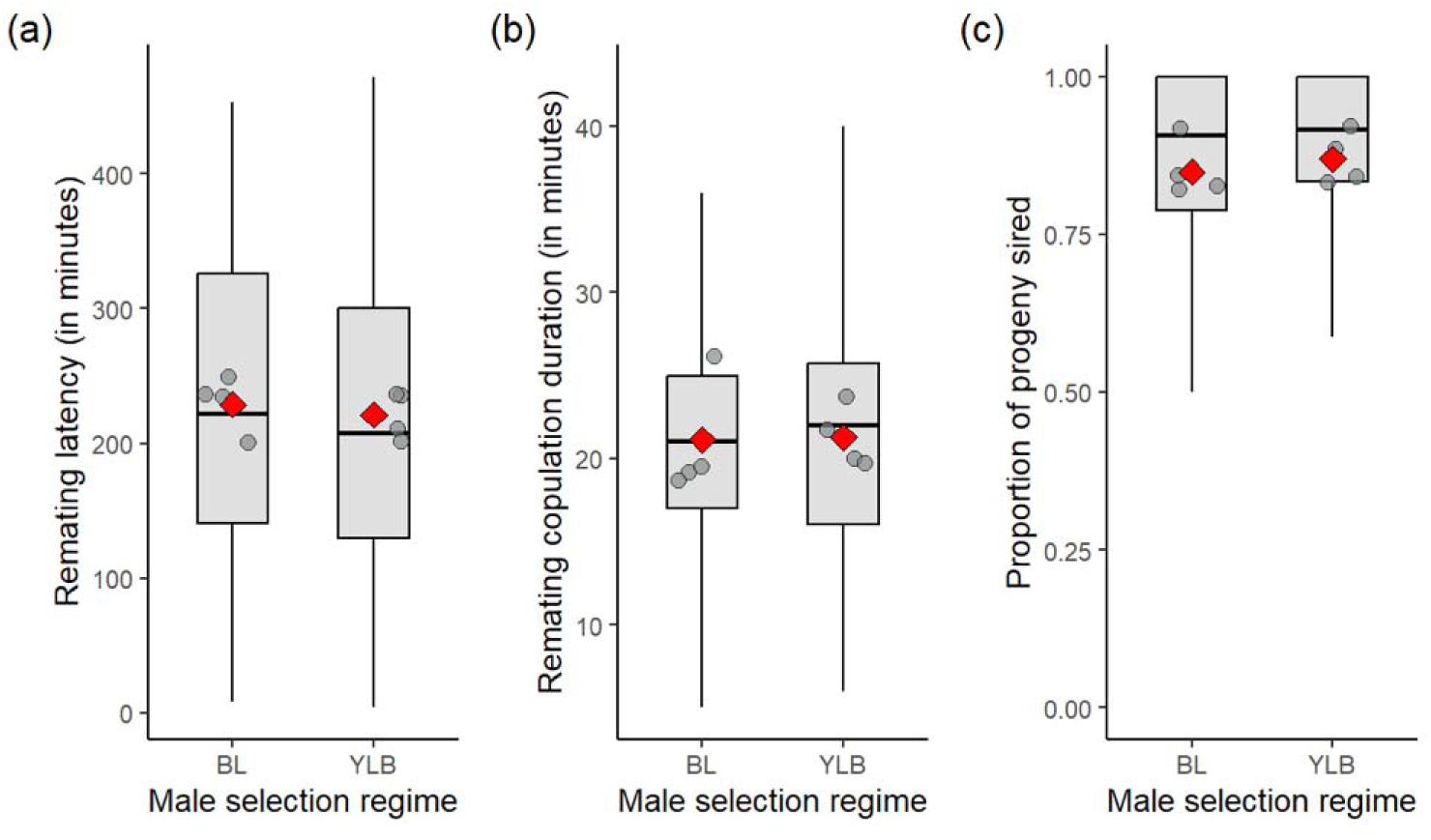
Boxplots showing (a) Remating latency, (b) Remating copulation duration and (c) P2 (proportion of progeny sired by target males) in experiment 3 (sperm offense assay). The grey dots represent the mean values of the replicate blocks and red dot represents to the overall mean across replicate blocks.

## Discussion

Our results demonstrate that YLB males do not differ significantly from BL control males across any of the evaluated fitness components. Consequently, chronic selection under adult live-yeast deprivation produced neither a direct nor a correlated evolutionary response in male fitness traits. Adaptation to this form of adult protein restriction appears to be strictly female-specific. Although we previously demonstrated marked female reproductive adaptation in these populations, specifically via increased early-life fecundity and altered ovarian functionality (Dasgupta et al., 2022), current findings clearly indicate that the underlying evolved female traits are decoupled from male fitness indicating a lack of intersexual genetic correlation (IGC) in the traits involved.

Sexual selection in males operates through both precopulatory and postcopulatory processes (Andersson, 1994; Birkhead & Møller, 1998). Precopulatory sexual selection relies heavily on male–male competition and energy-intensive courtship displays (Andersson, 1994). These traits are energetically costly and rely predominantly on dietary carbohydrate intake, as carbohydrates are rapidly metabolized to fuel neuromuscular activity (Lee et al., 2008; Maklakov et al., 2008; Jensen et al., 2015). In contrast, postcopulatory sexual selection including sperm competitive ability (both defense and offense) depends strongly on dietary protein (Simmons, 2001; Wigby et al., 2009). Beyond producing large quantities of sperm, males must synthesize a complex cocktail of seminal fluid proteins (SFPs) to form a functional ejaculate (Avila et al., 2011; McGraw et al., 2015). In Diptera, including *Drosophila melanogaster*, these proteins are primarily synthesized and secreted by the male accessory glands (Wolfner, 2002; Wigby et al., 2009). Accessory gland proteins (Acps) modulate female post-mating physiology by stimulating egg production, reducing female remating propensity, and maximizing male paternity share under polyandrous conditions (Chapman et al.,2003; Wolfner, 2002). Furthermore, Acps are essential for sperm storage, motility, and survival within the female reproductive tract (Neubaum & Wolfner, 1999; Poiani, 2006). Because the production of Acps is resource-intensive, postcopulatory trait expression is sensitive to male protein intake (Fricke et al., 2008). Ample empirical evidence highlights the role of the nutritional environment in modulating male postcopulatory traits. For example, McGraw et al. (2007) showed that *D. melanogaster* males reared on high-protein larval diets transferred more sperm, enhanced female sperm storage, and induced stronger remating refractoriness in their mates. Similarly, Amitin and Pitnick (2007) demonstrated that varying larval density and dietary yeast availability directly altered male sperm offense ability in *D. melanogaster*. Furthermore, adult protein intake can influence male reproductive success by altering courtship dynamics (Fricke et al., 2008; Duxbury & Chapman, 2020). Nevertheless, despite over 140 generations of selection under adult yeast deprivation, we observed no divergence in pre- or postcopulatory male competitive traits between evolved and ancestral populations. These results suggest that the role of adult diet in shaping male fitness is highly context-dependent and that selection under female-limiting yeast deprivation does not necessarily compromise or alter male reproductive performance.

Frameworks integrating nutrient-dependent life history trade-offs, such as the Geometric Framework for Nutrition, have consistently demonstrated distinct nutritional optima for male and female fitness (Maklakov et al., 2008; Rapkin et al., 2017; Hawkes et al., 2022). Female fitness is largely protein-constrained due to the massive nitrogenous investment required for egg production and vitellogenesis, whereas male fitness is primarily carbohydrate-dependent due to the high metabolic demand of courtship behaviours (Maklakov et al., 2008; Jensen et al., 2015). Under the chronic protein restriction imposed in our experimental regime, selection acted predominantly on female metabolic pathways to enhance the efficiency of resource provisioning for egg production.

The absence of IGC between male and female fitness traits in this system is notable, as theories of intralocus sexual conflict often predict shared genetic architecture to constrain sex-specific adaptation (Bonduriansky & Chenoweth, 2009; Schenkel et al., 2018). Males and females share highly conserved nutrient-sensing and metabolic signalling networks, including the insulin/insulin-like growth factor signalling (IIS) and target of rapamycin (TOR) pathways (Camus et al., 2019; Bennett-Keki et al., 2023). However, sex-specific regulation of these pathways allows for differential resource allocation: females prioritize resources toward oogenesis and vitellogenesis, whereas males prioritize somatic maintenance and energy mobilization (Mank, 2017; Han & Dingemanse, 2017). Furthermore, genes regulating nutrient acquisition, transport, and metabolism frequently exhibit sex-biased gene expression (Innocenti & Morrow, 2010; Mank, 2017). Such sex-specific transcriptional regulation reduces intersexual genetic correlation for fitness, thereby mitigating intralocus sexual conflict and enabling females to adaptively respond to chronic protein stress without imposing correlated fitness costs or benefits on males.

In summary, adaptation to chronic protein deprivation in YLB populations occurred through the modification of female-specific nutrient allocation and reproductive pathways, yielding a fully sex-limited evolutionary response unconstrained by intersexual genetic correlation.

## Acknowledgements

The experiments reported here was supported by the Core Research Grant (file no. CRG/2019/002460) from Science and Engineering Research Board, Department of Science and Technology, Government of India. SA thanks Department of Science and Technology, Government of India for financial support in the form of DST-INSPIRE Fellowship. We also thank Purbasha Dasgupta of Brown University for establishing the populations, and working on the initial version of the study that motivated a more complete set of experiments.

## Author contributions

BN conceived the idea, designed the experiment, analysed and interpreted the data, edited the manuscript, and secured funding. SA designed and performed the experiments, analysed data, and prepared the first draft of the manuscript. RSP, and SH helped in performing various assays and collected data.

