## Supplementary Information file for "No evidence of evolution of male fitness components in response to yeast deprivation: Putative female specific nutrient dependent reproductive adaptation"

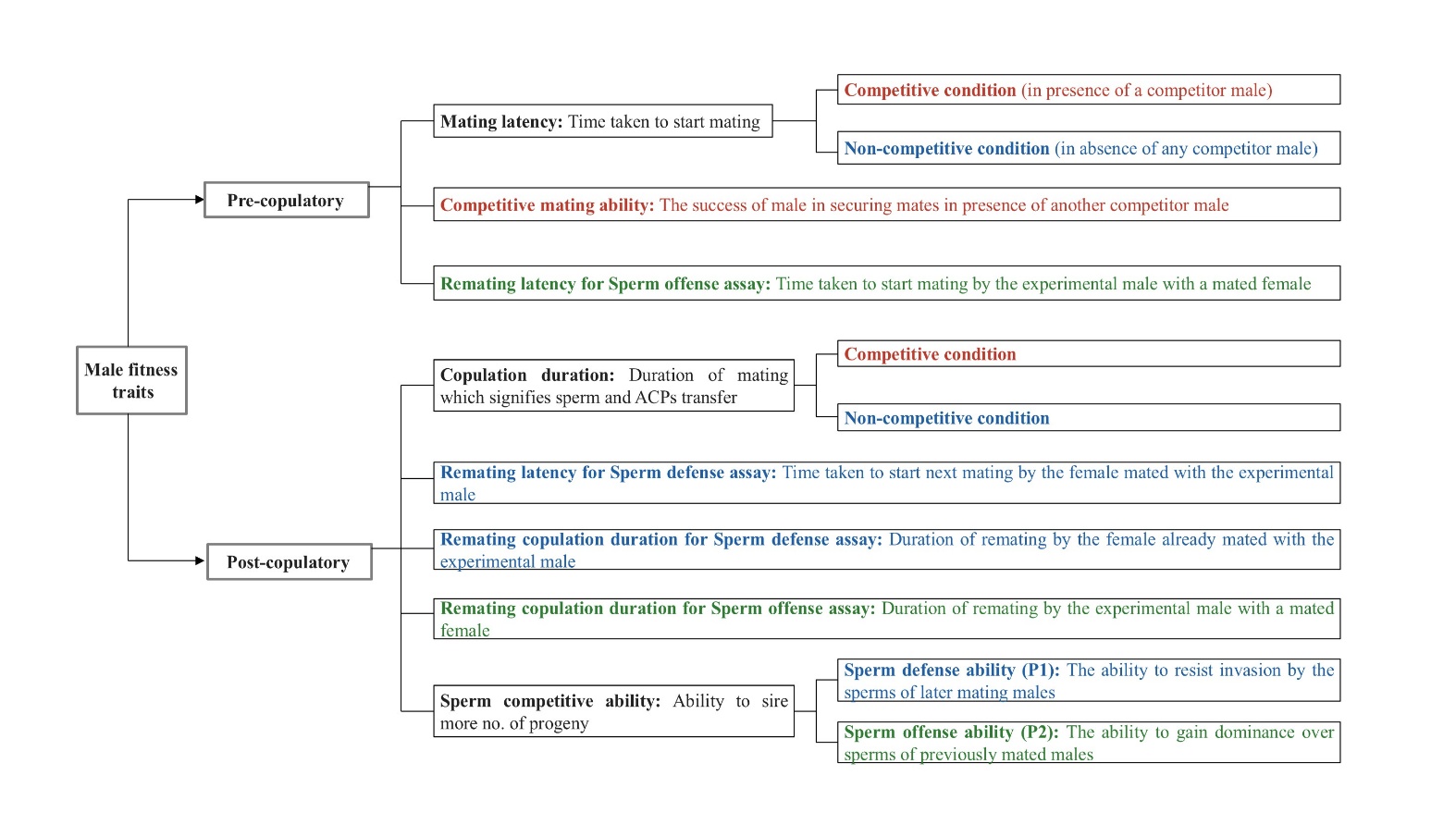

**Figure S1**: Flowchart of the male fitness traits measured in this study. Traits written in ‘red’ colour were measured in experiment 1 (Mating behaviour in presence of a competitor), those in ‘blue’ colour were measured in experiment 2 (Sperm defense assay), and ‘green’ colour were measured in experiment 3 (Sperm offense assay).

**Table S1: List of statistical models used for all the measured parameters.**

| **Experiment** | **Parameter** | **Model** |
| --- | --- | --- |
| **Experiment 1** | **Mating Success** | glmer(Mated_male ~ Regime+(1\|Block) +(1\|Block:Regime),family = binomial(link = "logit") |
|  | **Mating latency** | glmmTMB(ML~ Male_regime + (1\|Block) +(1\|Block:Male_regime), family = Gamma(link = "log") |
|  | **Copulation duration** | lmer(CD~Male_regime+(1\|Block)+(1\|Block:Male_regime) |
| **Experiment 2** | **Mating latency** | glmmTMB(ML~Male_regime + (1 \| Block) +(1\|Block:Male_regime), family = Gamma(link = "log") |
|  | **Copulation duration** | lmer(CD~Male_regime+(1\|Block)+(1\|Block:Male_regime) |
|  | **Mating fidelity (remating latency)** | glmer(RML ~ Male_regime + (1\|Block)+ (1\|Block:Male_regime), family = Gamma(link = "log") |
|  | **Remating copulation duration** | lmer(RCD~Male_regime+(1\|Block)+ (1\|Block:Male_regime) |
|  | **P1** | glmmTMB(cbind(red_eyed, Total-red_eyed) ~ Male_regime +(1\|Block)+ (1\|Block:Male_regime),  family = beta binomial(link="logit") |
| **Experiment 3** | **Remating latency** | lmer(RML~Male _regime+(1\|Block)+(1\|Block:Regime) |
|  | **Copulation duration** | lmer(RCD~Regime+(1\|Block)+(1\|Block:Regime) |
|  | **P2** | glmmTMB(cbind(red_eyed, Total-red_eyed) ~ Male_regime +(1\|Block)+(1\|Block:Male_regime), family = betabinomial(link="logit") |

**Table S2:** Block-wise results of effect of selection regime using generalized linear model (GLM) analyses of Mating latency under non-competitive condition in the experiment 2 (sperm defense assay). Considering α=0.05, significant p-values are highlighted in boldface.

| **Block** | **Sum sq.** | **Den. DF** | **Num. DF** | **F value** | **Pr(>F)** |
| --- | --- | --- | --- | --- | --- |
| BLOCK1 | 10.21 | 171 | 1 | 3.5387 | 0.06165 |
| BLOCK2 | 5.733 | 173 | 1 | 5.0564 | **0.0258** |
| BLOCK3 | 14.23 | 167 | 1 | 6.7842 | **0.01003** |
| BLOCK4 | 7.51 | 157 | 1 | 4.2978 | **0.0398** |

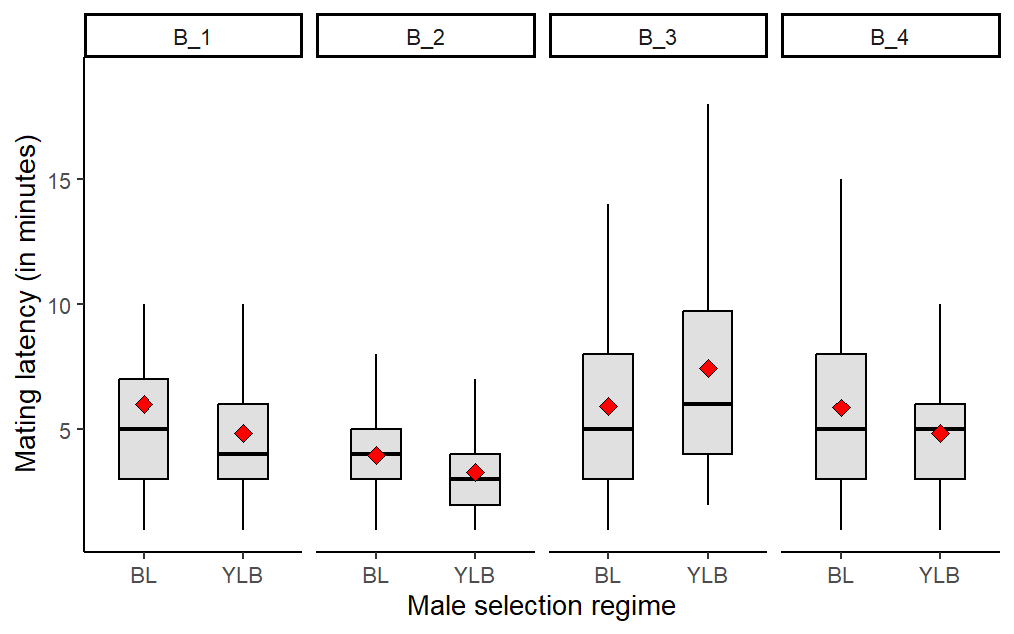

**Figure S2:** Boxplot showing the blockwise mating latency under non-competitive condition in the experiment 2 (sperm defense assay). The red dot represents to the mean value of the male selection regime.

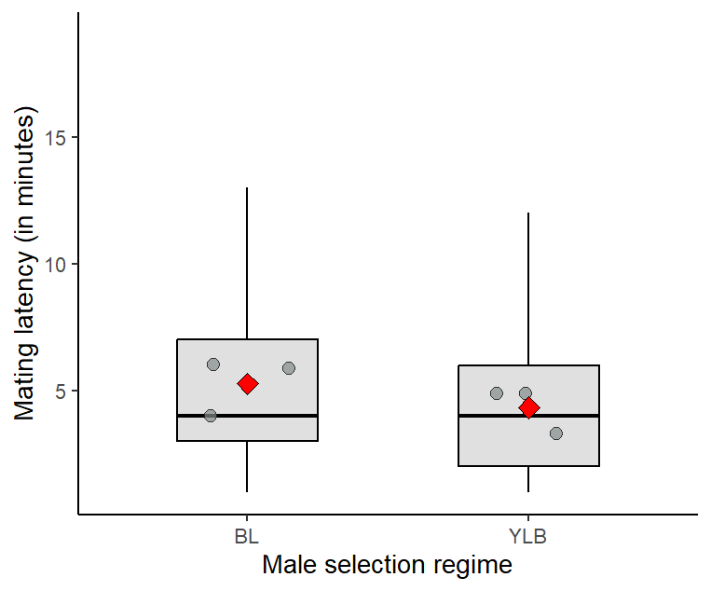

**Figure S3:** Boxplot showing mating latency under non-competitive condition in the experiment 2 (sperm defense assay) pooled across replicate blocks -1,2 and 4. The grey dots represent the mean values of the replicate blocks and red dot represents to the overall mean across replicate blocks. The effect of selection regime was tested with a generalized linear mixed model (GLMM) and was significant ($\chi^{2}$= 13.42, df=1, **p<0.05** ; α = 0.05**)**

**Table S3:** Block-wise results of effect of selection regime using generalized linear model (GLM) analyses of Copulation duration under non-competitive condition in the experiment 2 (sperm defense assay), considering α=0.05. Significant p-values are highlighted in boldface.

| **Block** | **Sum sq.** | **Den. DF** | **Num. DF** | **F value** | **Pr(>F)** |
| --- | --- | --- | --- | --- | --- |
| BLOCK1 | 0.001 | 189 | 1 | 0.001 | 0.972 |
| BLOCK2 | 88.57 | 168 | 1 | 7.436 | **0.007** |
| BLOCK3 | 0.657 | 175 | 1 | 2.329 | 0.129 |
| BLOCK4 | 211.72 | 169 | 1 | 14.245 | **<0.001** |

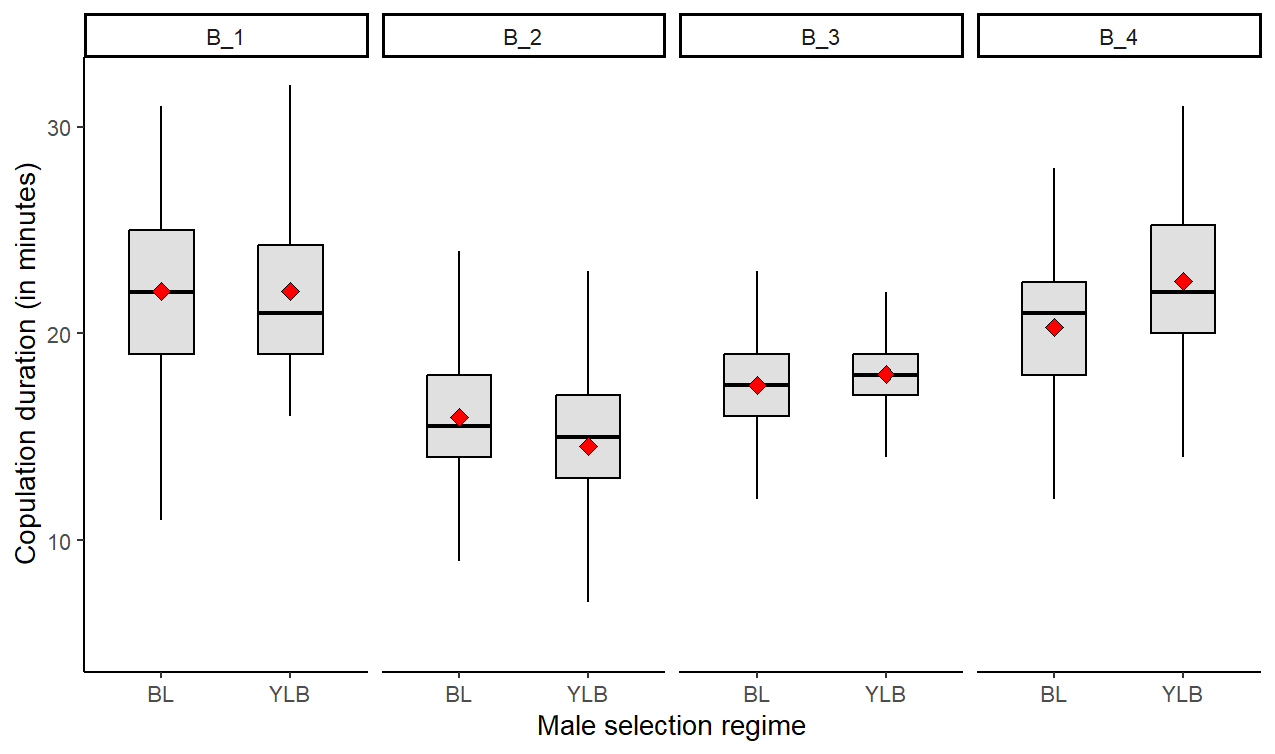

**Figure S4:** Boxplot showing the blockwise copulation duration under non-competitive condition in the experiment 2 (sperm defense assay). The red dot represents to the mean value of the male selection regime.

**Table S4:** Summary of the likelihood-ratio tests for the effect of the random factors- Block and Block x Male regime interaction on the measured traits of the experiment 1 (Mating behaviour). For each trait, models with and without the term of interest were compared using a χ² test. npar = number of estimated parameters; AIC = Akaike information criterion; LogLik = log-likelihood; DF = degrees of freedom. Significant p-values are in bold.

| **Traits** | **Effect** | **npar** | **AIC** | **LogLik** | **Chisq** | **DF** | **Pr(>Chisq)** |
| --- | --- | --- | --- | --- | --- | --- | --- |
| Mating latency | Block × Male regime | 5 | 1441.4 | -715.72 | 0.68 | 1 | 0.40 |
|  | Block | 4 | 1440.1 | -716.06 | 72.63 | 1 | **<0.05** |
| Copulation duration | Block × Male regime | 5 | 1860.5 | -925.26 | 0 | 1 | 1 |
|  | Block | 4 | 1858.5 | -925.26 | 33.40 | 1 | **<0.05** |
| Proportion of mating success | Block × Male regime | 4 | 891.32 | -441.66 | 0 | 1 | 1 |
|  | Block | 3 | 889.32 | -441.66 | 0.83 | 1 | 0.36 |

**Table S5:** Summary of the likelihood-ratio tests for the effect of the random factors- Block and Block x Male regime interaction on the measured traits of the experiment 2 (Sperm defense assay). For each trait, models with and without the term of interest were compared using a χ² test. npar = number of estimated parameters; AIC = Akaike information criterion; LogLik = log-likelihood; DF = degrees of freedom. Significant p-values are in bold.

| **Traits** | **Effect** | **npar** | **AIC** | **LogLik** | **Chisq** | **DF** | **Pr(>Chisq)** |
| --- | --- | --- | --- | --- | --- | --- | --- |
| Mating latency | Block × Male regime | 5 | 3222.7 | -1606.3 | 7.24 | 1 | **<0.05** |
|  | Block | 4 | 3227.9 | -1610 | 74.88 | 1 | **<0.05** |
| Copulation duration | Block × Male regime | 5 | 3817.6 | -1903.8 | 12.69 | 1 | **<0.05** |
|  | Block | 4 | 3828.2 | -1910.1 | 319.47 | 1 | **<0.05** |
| Mating fidelity (Remating latency) | Block × Male regime | 5 | 4989 | -2489.5 | 0 | 1 | 1 |
|  | Block | 4 | 4987.0 | -2489.5 | 5.68 | 1 | **0.01** |
| Remating copulation duration | Block × Male regime | 5 | 2319.1 | -1154.5 | 0 | 1 | 1 |
|  | Block | 4 | 2317.1 | -1154.5 | 3.20 | 1 | 0.07 |
| P1 | Block × Male regime | 5 | 1293.8 | -641.93 | 0 | 1 | 1 |
|  | Block | 4 | 1291.8 | -641.93 | 0 | 1 | 1 |

**Table S6:** Summary of the likelihood-ratio tests for the effect of the random factors- Block and Block x Male regime interaction on the measured traits of the experiment 3 (Sperm offense assay). For each trait, models with and without the term of interest were compared using a χ² test. npar = number of estimated parameters; AIC = Akaike information criterion; LogLik = log-likelihood; DF = degrees of freedom. Significant p-values are in bold.

| **Traits** | **Effect** | **npar** | **AIC** | **LogLik** | **Chisq** | **DF** | **Pr(>Chisq)** |
| --- | --- | --- | --- | --- | --- | --- | --- |
| Remating latency | Block × Male regime | 5 | 4008.8 | -1999.4 | 0 | 1 | 1 |
|  | Block | 4 | 4006.8 | -1999.4 | 0.59 | 1 | 0.44 |
| Remating copulation duration | Block × Male regime | 5 | 2061.4 | -1025.7 | 0.61 | 1 | 0.43 |
|  | Block | 4 | 2060.0 | -1026.0 | 27.639 | 1 | **<0.05** |
| P2 | Block × Male regime | 5 | 1212.6 | -601.28 | 0 | 1 | 1 |
|  | Block | 4 | 1210.6 | -601.28 | 287.77 | 1 | **<0.05** |

**Table S7:** Summary of the likelihood-ratio tests for the effect of the random factors- Block and Block x Male regime interaction on the mating latency measured in experiment 2 (Sperm defense assay) pooled across three replicate blocks- 1,2 and 4. Models with and without the term of interest were compared using a χ² test. npar = number of estimated parameters; AIC = Akaike information criterion; LogLik = log-likelihood; DF = degrees of freedom. Significant p-values are in bold.

| **Effect** | **npar** | **AIC** | **LogLik** | **Chisq** | **DF** | **Pr(>Chisq)** |
| --- | --- | --- | --- | --- | --- | --- |
| Block × Male regime | 5 | 2327.5 | -1158.8 | 0 | 1 | 1 |
| Block | 4 | 2325.5 | -1158.8 | 36.09 | 1 | **<0.05** |
